# U1 snRNP and RNA polymerase II interaction is predominantly mediated by Prp40 rather than U1-70K in yeast

**DOI:** 10.1101/2025.08.28.672894

**Authors:** Xueni Li, Jiaqin Li, Pankaj Srivastava, Shasha Shi, Aaron Issaian, Shaun Bevers, Marisa Elise Wagner, Angelo D’ Alessandro, Rui Zhao

## Abstract

Transcription and splicing are coupled both temporally and physically. A previous cryo-EM structure of the human U1 snRNP and RNA polymerase pol II complex has shown that U1 snRNP uses predominantly the RRM domain of U1-70K to directly interact with the RPB2 subunit of pol II. However, residues on U1-70K involved in the interaction with pol II are not conserved in yeast U1-70K, raising the question whether yeast U1 snRNP interacts with pol II in a similar manner. We found that yeast pol II associates with both U1 and U2 snRNPs, but U1-70K makes a minimal contribution to U1 snRNP’s interaction with pol II. On the other hand, multiple domains of yeast Prp40 interact with pol II and the removal of the C-terminal domain (CTD) of pol II does not affect this interaction. Although yeast Prp40 is stably associated with U1 snRNP, its human homologs, PRPF40a and PRPF40b, are alternative splicing factors that are not integral components of U1 snRNP. This shift of function of Prp40 homologs may have led to the evolution of U1-70K to be the main interactor with pol II in the human system.

## Introduction

In eukaryotes, DNA is first transcribed into pre-mRNA, which undergoes multiple processing events including splicing, before becoming mature mRNA. The two critical events in this process, transcription and splicing, are coupled both temporally and spatially. For example, a high percentage of splicing occurs co-transcriptionally and the speed of transcription can regulate splicing^1^. The transcription and splicing machinery also interact physically. RNA polymerase II (pol II), the main eukaryotic transcription machinery for protein-coding genes, contains 12 subunits, RPB1-12. The largest subunit, RPB1, contains a C-terminal domain (CTD) with multiple heptapeptide (YSPTSPS) repeats, 52 in human and 26 in yeast, that are heavily regulated by phosphorylation ^2^. The splicing machinery, the spliceosome, contains 5 snRNPs (U1, U2, U4, U5, and U6) and numerous non-snRNP factors ^3^. U1 snRNP is responsible for the initial recognition of the 5’ splice site (ss) and plays a key role in mediating the coupling between splicing and transcription. For example, U1 snRNP has been observed to associate with pol II ^4,5^, although it was unclear for a long time whether this interaction is direct. In 2021, Dr. Patrick Cramer’s lab determined the cryo-EM structure of human pol II and U1 snRNP assembled on a DNA-RNA scaffold made of a DNA mismatch bubble and a pre-mRNA containing the 5’ ss ^6^. This structure reveals that the RRM domain of U1-70K directly interacts with the RPB2 subunit of pol II. However, none of the human U1-70K residues that interact with RPB2 are conserved in yeast, making it unclear whether yeast U1 snRNP also interacts with pol II in a similar manner.

Yeast U1 snRNP is much larger and more complex than human U1 snRNP ^7^. The human U1 snRNP is composed of a 164-nt U1 snRNA, seven Sm proteins, U1A, U1C, and U1-70K proteins. The yeast U1 snRNP contains a much larger U1 snRNA (568 nt) and homologs of all the human U1 snRNP proteins (referred to as the U1 snRNP core). The yeast U1 snRNP core resembles the entire human U1 snRNP, with seven additional stably associated auxiliary proteins (Prp40, Luc7, Snu71, Prp39, Prp42, Snu56, Nam8) ^7^. It has been previously reported that yeast Prp40 interacts with pol II ^4^. It is not clear whether U1-70K and Prp40 both interact with pol II directly and what their relative contributions are to this interaction. In this paper, we set out to answer these questions using mostly biochemical approaches.

## Results

### Yeast pol II associates with both U1 and U2 snRNP

To evaluate the interaction between yeast pol II and U1 snRNP, we pulled down pol II from yeast whole cell lysate carrying the WT or CTD truncated pol II (with 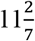or 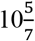heptapeptide repeats remaining) using the tandem affinity tag (TAP, protein A-calmodulin binding peptide (CBP)) tag on pol II subunit RPB3 ^8^. We then detected the snRNAs associated with pol II using solution hybridization ^9^ with fluorescent probes targeting all 5 snRNAs (U1, U2, U4, U5, and U6). We showed that both U1 and U2 snRNAs associate with pol II (**Fig. 1**), suggesting that yeast pol II interacts with both U1 and U2 snRNPs.

**Figure 1.**
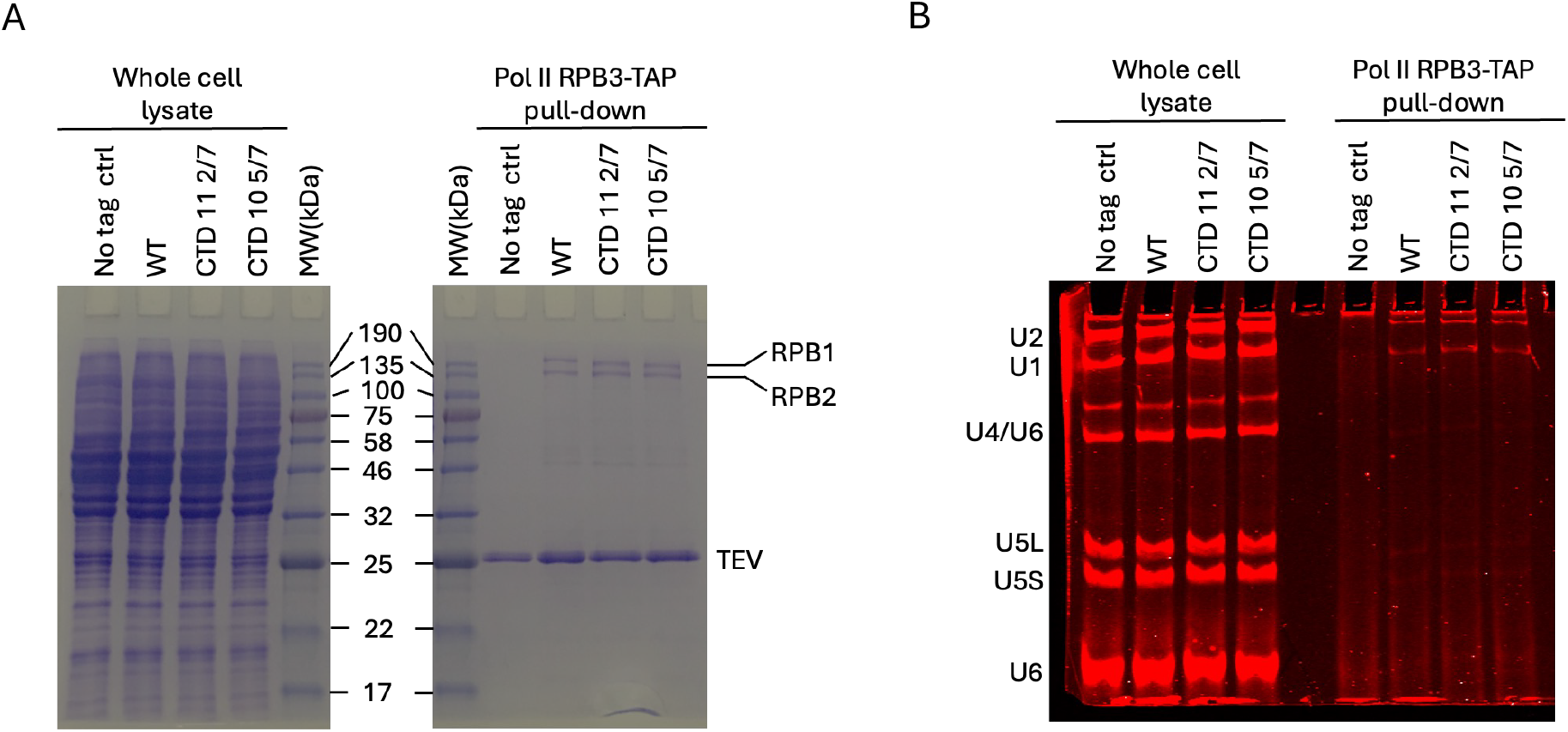
Yeast pol II associates with both U1 and U2 snRNPs. A.pol II WT or CTD truncations (11 2/7 or 10 5/7 heptapeptide repeats remaining) were pulled down from yeast cell lysate on IgG resin using the TAP tag on its RPB3 subunit and visualized on Coomassie-stained SDS PAGE. B.The pol II pulldown samples were analyzed for its snRNA content using solution hybridization and fluorescently labeled probes for each snRNA.

### U1-70K mutations and truncations do not substantially reduce pol II and U1 snRNP interaction in pull-down experiments

The cryo-EM structure of the human pol II and U1 snRNP complex showed that U1 snRNP predominantly uses a helix in the RRM domain of U1-70K to interact with RPB2 of pol II ^6^. However, the specific amino acids on U1-70K involved in the interaction are not conserved in yeast U1-70K (**Fig. 2A**). To evaluate whether any of the equivalent residues in yeast still contribute to pol II binding despite the lack of conservation, we generated five single amino acid mutations on U1-70K (E122K, K125E, K129E, F130A, K156E) in a U1-70K shuffle strain in which RPB3 of pol II was TAP-tagged. We pulled down pol II using IgG resin through the TAP tag on RPB3 and showed that pol II was associated with similar amounts of U1 snRNA from yeast strains expressing the mutant or the WT U1-70K (**Fig. 2B**).

**Figure 2.**
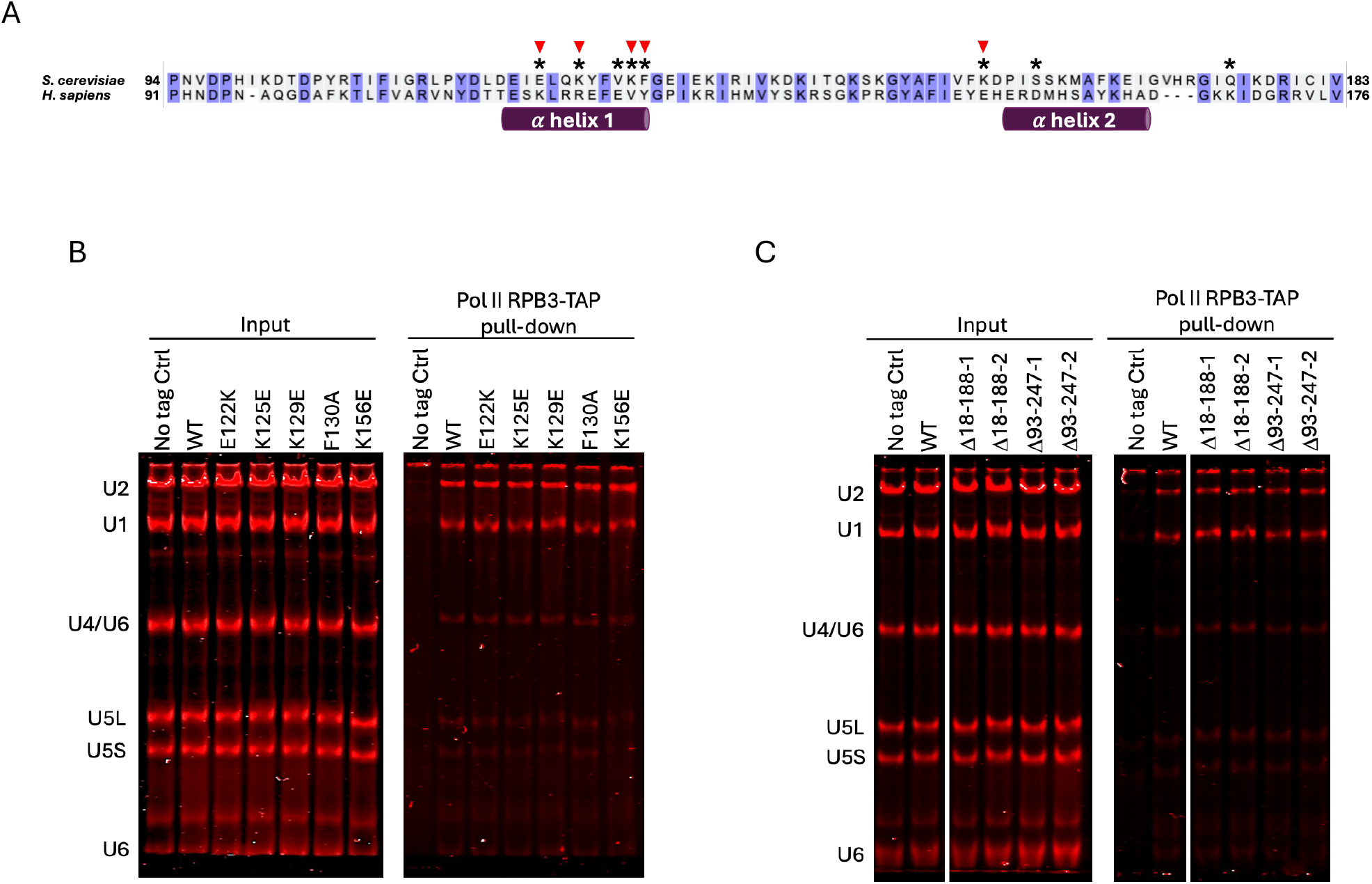
U1-70K mutants and truncations do not substantially reduce pol II and U1 snRNP interaction in pull down experiments from cell lysates. A.Amino acid sequence of the region in human U1-70K RRM domain that interacts with pol II in the cryo-EM structure of U1 snRNP and pol II complex ^6^ and its sequence alignment with the corresponding region in yeast U1-70K, using the alignment tool in Uniprot ^23^. Residues critical for the human U1 snRNP and pol II interaction based on the cryo-EM structure are labeled with*. Residues mutated in subsequent experiments are labeled with red triangles. B.pol II was pulled down from yeast strains carrying RPB3-TAP and various single amino acid mutations of yeast U1-70K residues corresponding to critical pol II-interacting residues in human U1-70K. The pull-down was analyzed for its snRNA content using solution hybridization and fluorescently labeled probes for each snRNA. C.pol II was pulled down from yeast strains carrying RPB3-TAP and U1-70K with two RRM domain truncations (Δ18-188 just removes the RRM domain and Δ93-247 removes a larger region containing the RRM domain which is known to be tolerated by yeast ^10^. The pull-down was analyzed for its snRNA content using solution hybridization and fluorescently labeled probes for each snRNA.

It is possible that multiple-residue mutations or residues other than the above in the RRM domain of U1-70K interact with pol II. To test this possibility and evaluate the role of the RRM domain of U1-70K in its interaction with pol II, we generated several U1-70K deletion mutants.

Δ93-247 is a deletion containing the RRM domain and surrounding residues that has been shown to be tolerated by yeast ^10^ and Δ108-188 removes only the RRM domain. We showed that pol II pulled down similar amounts of U1 snRNA in yeast carrying these deletions compared to those with the WT U1-70K (**Fig. 2C**). These data suggest that the RRM domain of U1-70K is not a major contributor to the interaction between U1 snRNP and pol II in yeast.

### U1-70K RRM makes a minimal contribution to the interaction between purified pol II and U1 snRNP

In the above pulldown experiments from cell lysate, the U1 and pol II association we observed could be through their direct interactions, but some of this association may also be contributed by both complexes binding to pre-mRNA, which could potentially blunt the difference in direct U1 and pol II interaction elicited by U1-70K mutants. To rule out this possibility, we evaluated the direct interaction between pol II and U1 snRNP using purified complexes. We purified pol II using the TAP (CBP-TEV-protein A) tag on RPB3 and U1 snRNP using the SBP-TEV-protein A tag on U1A from yeast. Mass spectrometry analysis of the purified U1 snRNP sample shows it contains most of the pol II subunits (**Fig. 3A**), consistent with the interaction between U1 snRNP and pol II. We first used U1 snRNP WT or quintuple mutant (E122K/K125E/K129E/F130A/ K156E) on IgG resin to pull down purified pol II. We found that U1 mutant and WT pull down similar level of pol II (**Fig. 3B**). We next used pol II (containing CBP tag on RPB3) on calmodulin resin to pull down purified U1 snRNP containing either U1-70K WT or ΔRRM (residues 108-188 deleted). We found that pol II pulls down slightly less U1 snRNP ΔRRM compared to the WT (**Fig. 3C**), although removing the RRM is far from abolishing the interaction, as would be expected based on the human U1 snRNP and pol II cryo-EM structure ^6^.

**Figure 3.**
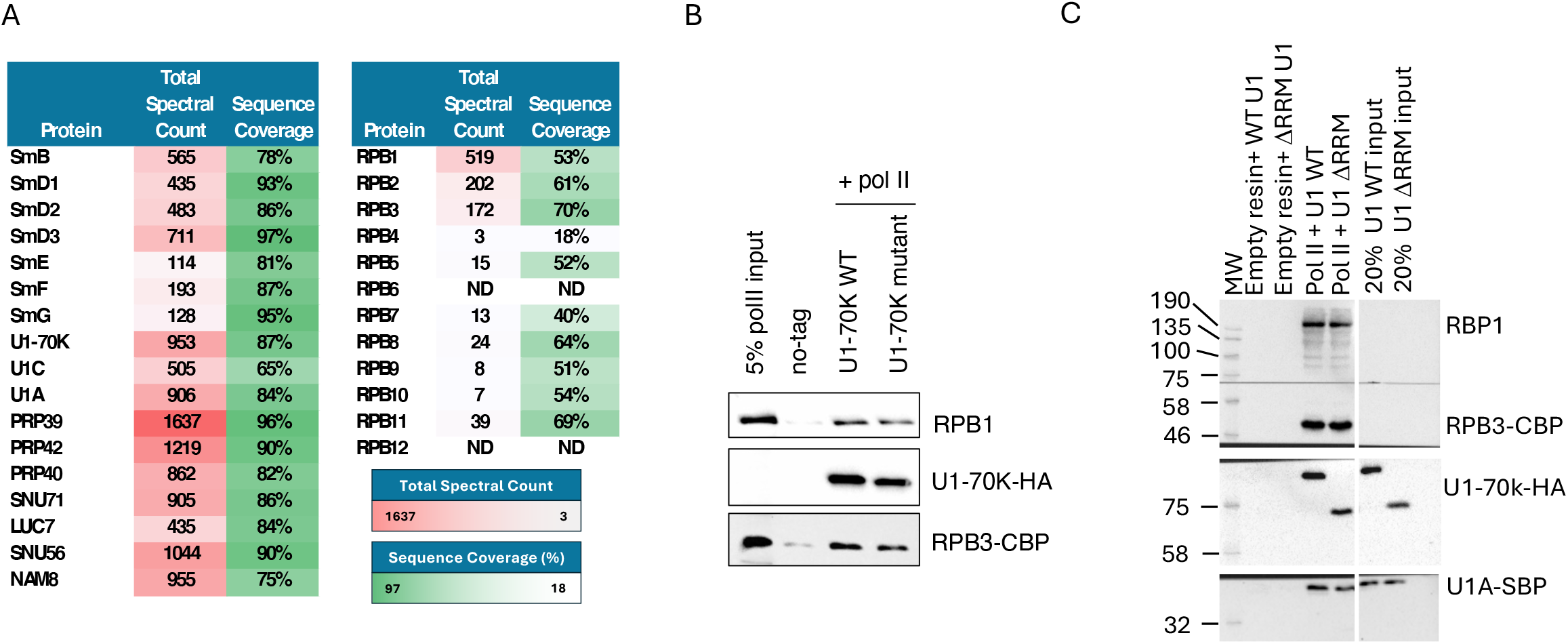
Deletion of the RRM domain in purified U1 snRNP has minimal impact on its interaction with purified pol II. A.Mass spectrometry analysis of the purified U1 snRNP sample shows it contains most of the pol II subunits. ND, not detected. B.U1 snRNP WT or quintuple mutant was immobilized on IgG resin and used to pull down purified pol II. The results were analyzed using Western blot, with U1-70K, RPB1, and RPB3 detected by an anti-HA, the anti-RPB1 antibody 8WG16, and anti-CBP antibody, respectively. C.Purified pol II (CBP-tagged) was immobilized on calmodulin resin and used to pull down purified U1 snRNP WT or ΔRRM, which was then analyzed by Western blot. U1-70K and U1A components of the U1 snRNP were detected using an anti-U1-70K antibody or an anti-SBP antibody. RPB1 and RPB3 were detected by the anti-RPB1 antibody 8WG16 and anti-CBP antibody, respectively.

### Prp40 makes a substantial contribution to the interaction between U1 snRNP and pol II

Given that U1-70K does not seem to be making major contributions to the interaction between U1-70K and pol II (**Fig. 2, 3**), while U1 snRNP and pol II clearly interact, another U1 snRNP component may be making a more important contribution to this interaction. It has been previously shown that Prp40 interacts with pol II ^4^. We therefore evaluated the interaction between Prp40 and pol II and compared it with the interaction between U1-70K and pol II.

To this end, we took advantage of a yeast strain carrying Luc7 with an inducible degron and TAP-tagged U1A (for U1 snRNP purification) that we had previously generated ^11^. We have previously shown that Luc7 depletion also removes Prp40 and Snu71 from U1 snRNP ^11^. We modified the CBP in the TAP tag with SBP, so that it does not carry the same CBP tag that is on RPB3 of pol II. We purified U1 snRNP without Prp40, Luc7, and Snu71 from this strain as well as U1 snRNP carrying U1-70K with the RRM domain deleted (**Supplemental Fig. S1**). We used purified pol II to pull down purified U1 snRNP WT, U1-70K ΔRRM, or ΔPrp40/Luc7/Snu71 and detect the amount of U1 snRNP in the pull down using an anti-SBP antibody. We showed that pol II pulls down slightly less U1 snRNP ΔRRM (**Fig. 4A**). On the other hand, pol II pulls down dramatically less U1 snRNP ΔPrp40/Luc7/Snu71 compared to the WT (**Fig. 4A**), indicating that Prp40/Luc7/Snu71 contributes more substantially to the interaction with pol II than U1-70K.

**Figure 4.**
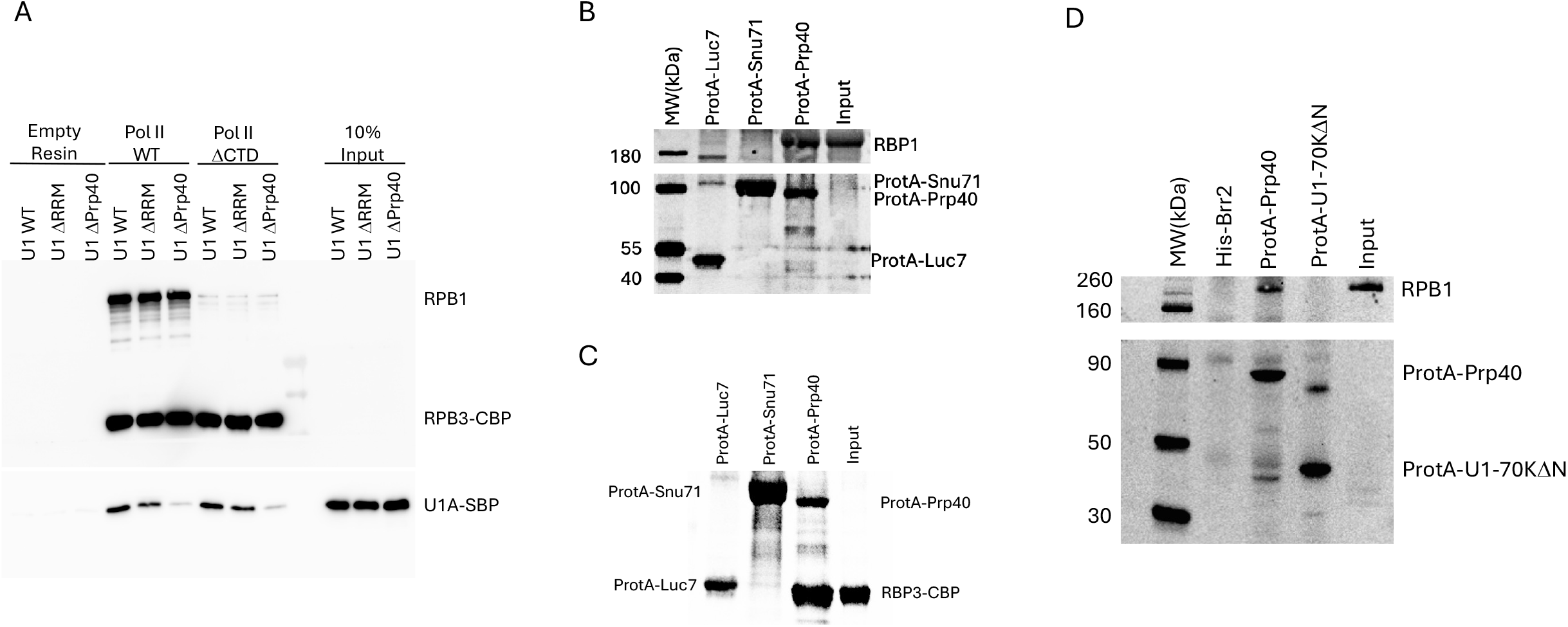
Pol II interacts with Prp40 much more strongly than with U1-70K, and CTD truncation does not affect these interactions. A.Purified pol II WT or ΔCTD (both were CBP tagged) was immobilized on calmodulin resin and used to pull down purified U1 snRNP WT or ΔRRM or ΔPrp40/Luc7/Snu71. Pol II was probed with both the anti-RPB1 antibody 8WG16 (which cannot recognize pol II ΔCTD) and an anti-CBP antibody (which recognizes RPB3-CBP). U1 snRNP was probed with an anti-SBP antibody that recognizes U1A-SBP. B.Purified protein A-tagged Prp40 or Snu71 or Luc7 were immobilized on IgG resin and used to pull down purified pol II, which was probed by the anti-RPB1 antibody 8WG16 in Western blot. The antibody also detected protein A-tagged Prp40, Snu71, or Luc7 through their protein A tag. C.The same as in panel B, but probed with an anti-CBP antibody against PRB3-CBP. The antibody also detected protein A-tagged Prp40, Snu71, or Luc7 through their protein A tag. D.Purified protein A-tagged Prp40 or U1-70K ΔN (residues 1-97 deleted) were immobilized on IgG resin and used to pull down purified pol II, which was probed by the anti-RPB1 antibody 8WG16 in Western blot.

To evaluate if the CTD of pol II is important for its interaction with Prp40/Luc7/Snu71, we inserted a PreScission site between residues 1513 and 1514, which are between the main body of RPB1 and its CTD. After purifying pol II, we cleaved off the CTD using PreScission protease and used the pol II ΔCTD (**Supplementary Fig. S1**) to pull down various U1 snRNPs. We showed that the pulldown pattern is essentially identical to that of WT pol II (**Fig. 4A**), indicating that the CTD does not play a major role in the interaction between pol II and Prp40/Luc7/Snu71 (as well as in the weak interaction between U1-70K and pol II). This result is consistent with our observation that using a TAP tag on RPB3 to pull down either WT or CTD-truncated pol II yields similar amounts of associated U1 snRNA (**Fig. 1**).

To determine if Prp40 alone in the Prp40/Luc7/Snu71 trimer is sufficient to interact with Pol II, we purified protein A-tagged Prp40 alone on IgG resin from yeast. We showed that it can efficiently pulldown purified pol II which is detected using the anti-RPB1 antibody 8WG16 (**Fig. 4B**) or an anti-CBP tag against RBP3-CBP (**Fig. 4C**). On the other hand, protein A-tagged Snu71 or Luc7 alone does not pull down any significant amount of pol II (**Fig. 4B, C**).

We further purified protein A-tagged U1-70K ΔN (residues 1-97 deleted but RRM domain remained intact, since we were unable to express full length U1-70K) from yeast and used it or protein A-tagged Prp40 to pull down purified pol II. We showed that purified Prp40 pulled down significantly more pol II compared to U1-70K ΔN (**Fig. 4C**), confirming our observation that Prp40 makes a more significant contribution to pol II interaction than U1-70K.

### Multiple domains of Prp40 interact with pol II

To decipher which domain of Prp40 interacts with pol II, we expressed and purified GST-fused WW, FF1, 2, 3, 4, 5, 6, and 6 extension domains (**Fig. 5A**) in E. coli and used them to pull down purified pol II. We found that the FF4 domain has the strongest interaction, followed by FF2, FF1, FF3, and the extension downstream of FF6, while the WW, FF5, and FF6 domains have no substantial interactions (**Fig. 5B**).

**Figure 5.**
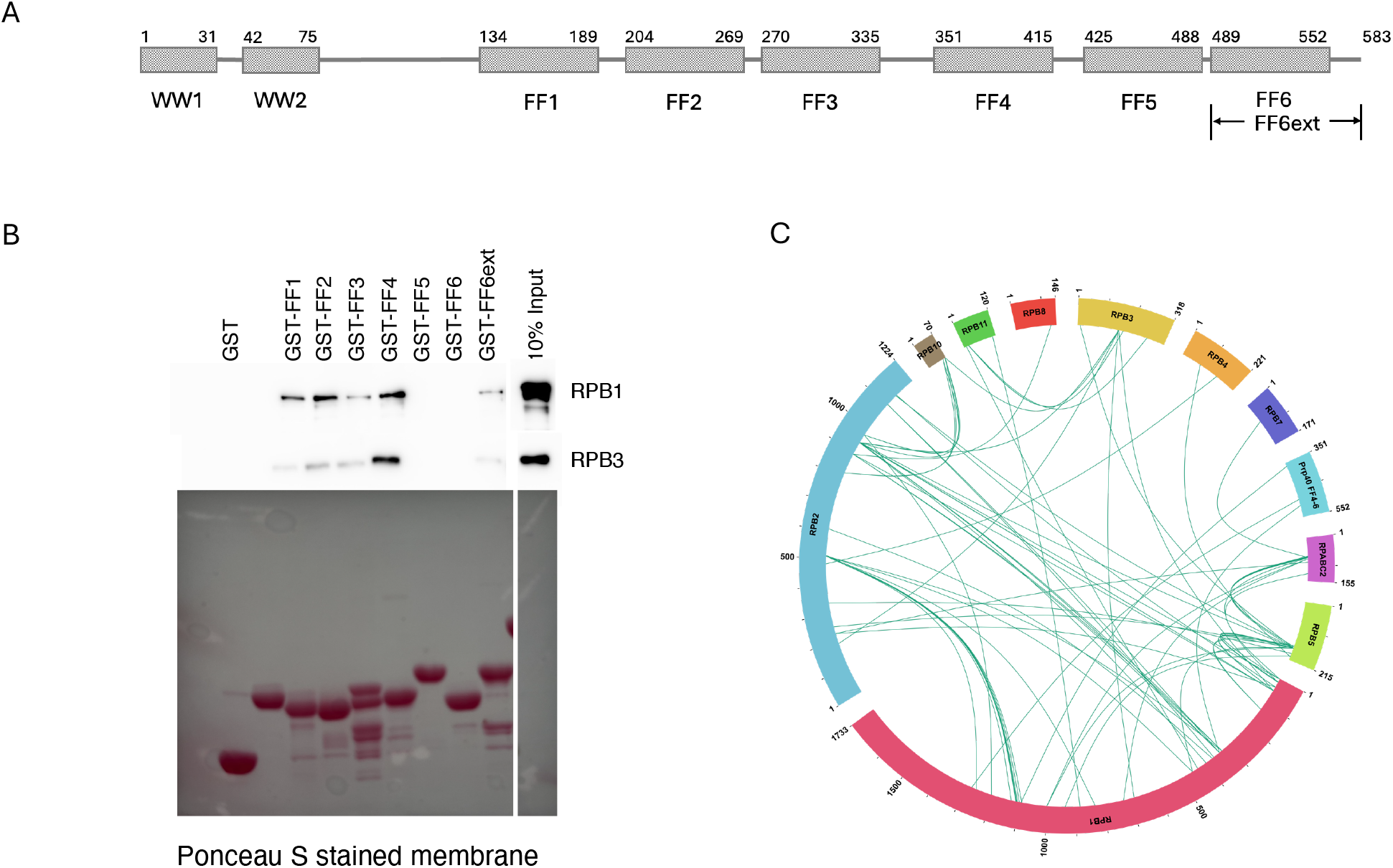
Multiple domains in Prp40 interact with pol II. A.A schematic diagram showing the domain organizations of Prp40. Residue numbers represent the boundary of the constructs used in this experiment. B.Purified GST-fused domains of Prp40 were immobilized on glutathione resin (shown on bottom SDS PAGE gel by ponceau S stain) and used to pull down purified pol II, which was probed by the anti-RPB1 antibody 8WG16 or anti-CBP antibody which recognizes RPB3-CBP in Western blot. C.Purified Prp40 FF4-6 is incubated with pol II, crosslinked with DSSO or SDA, and analyzed by mass spectrometry with the identified crosslinks from both crosslinkers displayed in a wheel diagram.

We next incubated purified pol II with the Prp40 FF4-6 domain and treated the sample with the DSSO or SDA crosslinker. Mass spectrometry analysis of the crosslinked sample revealed that the FF4-6 domains were crosslinked to RPB1 and 5 (**Fig. 5C**), supporting the interaction between the two proteins observed in pull-down analyses (**Fig. 5B**).

## Discussion

U1 snRNP plays an important role in the physical coupling between transcription and splicing, connecting the spliceosome to pol II. In humans, cryo-EM structural studies have shown that U1 snRNP uses its RRM domain to interact with RPB2 of pol II ^6^. However, residues on U1-70K that interact with RPB2 are not conserved in yeast U1-70K ^6^. In this paper, we showed that U1-70K does not play a major role in interacting with pol II in yeast (**Figs. 2-4**). Deletion of the RRM domain of U1-70K only minimally affected the interaction between purified U1 snRNP and pol II. On the other hand, Prp40 seems to be the main player mediating the interaction between U1 snRNP and pol II (**Fig. 4**). We found that multiple domains of Prp40 (including FF1-4 and FF6 extension) directly interact with purified pol II in pull-down assays using GST-Prp40 domains purified from E. coli, which was consistent with our crosslinking and mass spectrometry analyses (**Fig. 5**).

Prp40 has been previously reported to interact with pol II by Morris and colleagues, although this interaction was thought to be with the CTD of pol II ^4^. In Morris et al., GST-fused Prp40 purified from E coli was run on SDS PAGE, transferred to nitrocellulose membrane, and incubated with radiolabeled pol II CTD to detect binding in a Far Western assay. The GST-Prp40 proteins used in Far Western analyses were denatured on SDS-PAGE, which may generate artifacts in interaction studies. The GST-Prp40 FL and domains used in these studies also contained a substantial amount of contaminating proteins, potentially complicating the interpretation of the results. In our experiments, using highly purified pol II and U1 snRNP, we showed that deleting the entire CTD from pol II had no effect on the interaction between pol II and U1 (**Fig. 4A**). The lack of CTD involvement may help maintain a more stable interaction without it being influenced by the numerous regulatory events occurring on the CTD.

Our results revealed interesting differences in the physical coupling between U1 snRNP and pol II in yeast and human. In yeast U1 snRNP, Prp40 is a stably associated component, but its human homologs, PRPF40a and PRPF40b, are alternative splicing factors ^12-14^ that are no longer integral components of U1 snRNP. The lack of stable association of human PRPF40s with U1 snRNP may have led to the evolution of U1-70K to be the main interactor between U1 snRNP and pol II. This may also free up PRPF40s to function as alternative splicing factors without compromising the coupling between splicing and transcription. Indeed, multiple sequence alignment (**Fig. 6**) shows that most key residues in U1-70K that are important for pol II binding are highly conserved among higher eukaryotes with substantial alternative splicing, but are not conserved in *S. cerevisiae, S. pombe,* and *C. albicans*, which have few to no alternative splicing. This also correlates with a higher sequence similarity between *S. pombe* or *C. albicans* Prp40 and *S. cerevisiae* compared to those from higher eukaryotes (**Fig. 6B**).

**Figure 6.**
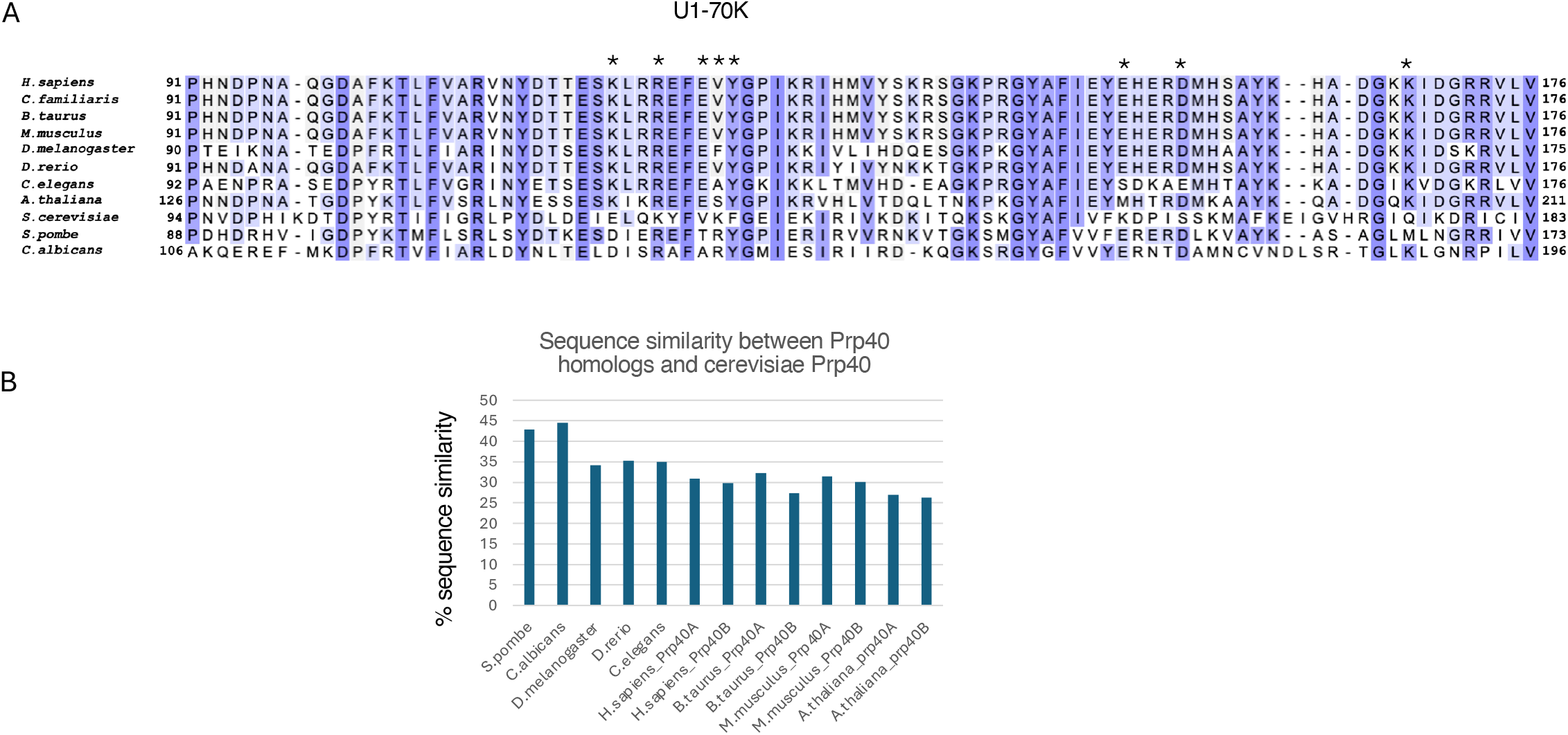
Sequence comparison of U1-70K or Prp40 homologs among different species. A.Multiple sequence alignment of the region of U1-70K involved in interaction with pol II among different species. Blue designates conserved residues (with darker colors indicating conservation among more species than the lighter colors). * designates residues important for the interaction with pol II based on the cryo-EM structure of human U1 snRNP and pol II ^6^. B.Sequence similarities between Prp40 homologs and S. cerevisiae Prp40. Sequence alignment was performed using the EMBL-EBI EMBOSS Needle Pairwise Sequence Alignment tool ^24^.

We also found that yeast pol II interacts with both U1 and U2 snRNP (**Fig. 1**). This is consistent with previous observations by Robert and colleagues who found that a complex containing RNA pol II purified using beads coupled to transcription elongation factor SII also contained U2 snRNP ^15^. The molecular details of the U2 snRNP and pol II interaction are not yet clear, although TAT-SF1, a U2 snRNP component, has been shown to interact with pol II ^16^.

The coupling between transcription and splicing is potentially important for efficient co-transcriptional splicing. Interestingly, splicing seems to also influence transcription, potentially through their coupling. For example, Caizzi and colleagues found that inhibition of pre-mRNA branch site recognition by U2 snRNP increases the duration of pol II pausing in the promoter-proximal region, hinders the recruitment of the pause release factor P-TEFb, and reduces pol II elongation velocity at the beginning of genes ^17^. Our investigation of the U1 snRNP and pol II interaction in yeast is a step towards better understanding the function of this coupling in the future.

## Materials and Methods

### pol II pull-down and snRNA detection

RPB3 TAP tagged yeast strains expressing RPB1 wild-type, CTD 11 2/7 and 10 5/7 truncations were gifts from Richard Young’s lab ^8^. The yeast cells were cultured to an OD600 of 3, harvested and lysed using bead-beating methods in buffer IgG 150 (20mM Tris 8.0, 150mM NaCl, 0.02% NP-40, 1mM DTT supplemented with protease inhibitors). The cleared cell lysate was incubated with IgG resin for 3 hr to pull down pol II. The resins were washed thoroughly, and pol II was cleaved off the resin by TEV protease. RNAs associated with pol II were released using proteinase K and analyzed by solution hybridization with IRDye700-labeled probes specific for U1, U2, U4, U5, U6 snRNAs ^9^.

To assess the effects of mutations on the RRM domain of yeast U1-70K (SNP1) mutations, we integrated a TAP tag to the C termini of RPB3 using *HIS3* marker in the *snp1Δ* [pRS316-SNP1] shuffle strain (gift from Stewart Shuman’s lab) ^18^. A 1.59-kbp DNA segment bearing the SNP1 gene (nucleotides -400 to +1190) was amplified from pRS413-SNP1 (gift from Stewart Shuman’s lab) ^18^ and inserted into pRS415 vector to make pRS415-SNP1. E122K, K125E, K129E, F130A, K156E single mutations or Δ108-188, Δ93-247 deletions were generated on pRS415-SNP1. The plasmids were transfected into the RPB3-TAP SNP1 shuffle strain. After FOA selection, the resulting strains were used to perform the same pol II pulldown and snRNA detection experiment as above.

### pol II purification

Yeast pol II was purified by using the TAP tag on RPB3 ^8^. Specifically, 6 liters of cells were cultured in YEPD medium at 30 °C to an OD600 of 4. The cell pellets were re-suspended in 10 ml of lysis buffer (50 mM Tris-HCl, pH 8.0, 200 mM NaCl, 0.05% NP-40, 1mM DTT). The cell suspension was snap-frozen into liquid nitrogen to form yeast “popcorn” and cryogenically ground using a SPEX 6870 Freezer/Mill. The frozen cell powder was thawed at room temperature and re-suspended in an additional 60 ml of lysis buffer with protease inhibitor cocktails (Sigma Aldrich). The cell lysate was first centrifuged at 27,845×g for 1 h in a GSA rotor (Sorvall) and the supernatant was further centrifuged at 167,424×g rpm in a 45Ti rotor (Beckman) for 1h at 4 °C. The supernatant was incubated with 2 ml of IgG Sepharose-6 Fast Flow resin (GE Healthcare) overnight at 4 °C. The resin was washed with IgG washing buffer (20 mM Tris-HCl, pH 8.0, 200 mM NaCl, 0.02% NP40, 0.5 mM dithiothreitol (DTT)) and then washed with the same buffer containing 500mM NaCl then 700mM NaCl. The resin was incubated with TEV protease in 1 ml TEV150 buffer (20 mM Tris-HCl, pH 8.0, 150 mM NaCl, 0.5 mM DTT) overnight at 4 °C. The purified pol II was slightly concentrated using an Amicon centrifugal filter, aliquoted and frozen until use.

To purify pol II with the entire CTD truncated, we inserted a PreScission cutting site into RPB1 before its CTD region (between residues 1513 and 1514) and the modified plasmid was transformed into RPB3 TAP-tagged yeast strains for the RPB1 shuffling experiment (strain GHY1729 ^8^). After FOA selection, the strain was cultured and pol II was purified in the same way, except for the addition of an incubation step of the resin with PreScission protease and then washes to remove the CTD region, before the final release of purified pol II using TEV protease.

### U1 snRNP purification

To purify wild-type and SNP1-RRM deleted U1 snRNP, we integrated a STP tag (SBP-TEV-protA) to the C terminus of U1A using *HIS3* marker in *snp1Δ* [pRS316-SNP1] strain, transformed pRS415-SNP1-2xHA, pRS415-SNP1(Δ108-188)-2xHA into it and shuffle out pRS316-SNP1 using FOA selection. Twelve liters of cells of each SNP1 construct were cultured in YEPD medium at 30 °C to an OD600 of 4 and harvested for U1 snRNP purification.

To purify PRP40-depleted U1 snRNP, we introduced an STP tag (SBP-TEV-protA) to the C-terminus of U1A using *HIS3* marker in *Luc7Δ* [pRS316-LUC7] strain ^19^. The plasmid bearing auxin-inducible degron (AID) plus 3x-HA tag at the C terminus of LUC7 and an auxin receptor (OsTIR1) driven by a β-estradiol inducible promoter ^11^ was transformed into the U1A STP-tagged *luc7* [pRS316-LUC7] strain. After FOA selection, the strain was cultured in YEPD medium at 30 °C to an OD600 of 1, and 1 ml each of β-estradiol (0.116g/50mL of ethanol) and IAA (6.58g/50ml of ethanol) were added per liter cell to induce the degradation of LUC7 for 4 hr. Twelve liters of cells were cultured and harvested for U1 snRNP purification. U1 snRNP purification was performed essentially the same way as previously described ^7^, but without the 2^nd^ purification using calmodulin resin.

### Pull-down assay to evaluate the interaction between pol II and U1 snRNP *in vitro*

Purified pol II and U1 snRNP were incubated at ∼1:1 molar ratio in a buffer containing 20 mM Tris-HCl, pH 8.0, 150 mM NaCl, 2mM MgCl2, 1mM CaCl2, 0.5 mM DTT for 2 hr at 4 °C. Pol II and associated U1 snRNP were pulled onto calmodulin resin, taking advantage of the CBP tag remained on pol II after purification. The resins were washed using the same buffer as above, and proteins were eluted using a buffer containing 20 mM Tris-HCl, pH 8.0, 150 mM NaCl, 2mM EGTA, 0.5 mM DTT. The eluted proteins were separated on SDS-PAGE and transferred to a nitrocellulose membrane. Western blots were performed to detect RPB1, RPB3-CBP, U1A-SBP or SNP1-2xHA. Antibodies used were RPB1 antibody (8WG16, ref), CBP tag antibody (Genscript), SBP tag antibody (Santa Cruz) and HA tag antibody (3F10, Roche).

### Pull-down assay to evaluate the interaction between pol II and U1-70K ΔN, PRP40, or SNU71/LUC7

The coding regions of yeast PRP40, SNU71, and LUC7 full length and N-terminal truncated SNP1 (SNP1 Δ1-97) were amplified by PCR using genomic S. cerevisiae DNA as a template, and ligated into pRS414, pRS416 and pRS317 vectors. The final plasmids constructed are: pRS414/GPD-protA-TEV-PRP40, pRS414/GDP-protA-TEV -SNU71, pRS317/GPD-LUC7 and pRS414/GPD protA-TEV-SNP1 Δ1-97. Yeast BCY123 cells transformed with pRS414/GPD-protA-TEV-PRP40 or pRS414/GPD protA-TEV-SNP1 Δ1-97 or co-transformed with pRS414/GDP-protA-TEV-SNU71 and pRS317/GPD-LUC7 were kept and cultured using appropriate selective media. One hundred to 500 ml of cells cultured to an OD600 of 3-4 were used for purification. Cells were harvested and lysed in lysis buffer (40 mM Tris-HCl, pH 8.0, 200 mM NaCl, 0.05% NP40, 1 mM DTT, protease inhibitors, 5ul Benzonase/ml) using the bead-beating method. The lysates were applied to IgG resin. The resins were washed with the lysis buffer and incubated with 0.4 μM purified pol II (TEV protease contains a His tag and was removed from pol II using Ni-NTA resin) in a buffer containing 20 mM Tris-HCl, pH 8.0, 150 mM NaCl, 0.5 mM DTT for 1.5 hr at 4 °C. The resins were washed, and the proteins were cleaved off the IgG resin using TEV protease in the same buffer. The proteins were separated on SDS-PAGE and transferred to a nitrocellulose membrane. Western blot was performed using RPB1 antibody and anti-CBP antibody as the above.

### GST pull-down assays using purified proteins

Various PRP40 domains fused with an N-terminal Flag tag and SNP1 RRM domain (residue 98-188) were cloned into the pGEX-6p-1 vector (GE Healthcare) and purified from E. coli as GST fusion proteins. The Prp40 domains constructed are: 1-75 (WW); 134–189 (FF1); 204-269 (FF2); 270–335 (FF3); 351-415 (FF4); 425–488 (FF5); 488-552 (FF6); 488-583 (FF6 ext) and 351–552 (FF4-6). To detect interactions between Prp40 domains with pol II, GST fusion proteins or GST alone were bound on glutathione-Sepharose resin (GE Healthcare) and incubated with 1μM of purified pol II for 1.5 hr at 4 °C. The resins were washed, and the proteins were cleaved off the glutathione-Sepharose resin using PreScission protease. The proteins were separated on SDS-PAGE and transferred to a nitrocellulose membrane. Western blot was performed using RPB1 antibody and CBP tag antibody as the above.

### Crosslinking and LC-MS analyses

Purified pol II and Prp40 FF4-6 protein were mixed at ∼1:1.5 molar ratio and dialyzed against buffer (20 mM HEPES pH 7.5, 150 mM NaCl, 1 mM MgCl_2_ and 0.5 mM TCEP). Samples were crosslinked with either 1 mM disuccinimidyl sulfoxide (DSSO, Thermo Fisher) or 2 mM NHS-Diazirine (SDA, AAT Bioquest) for 1 hour at 4 °C. Residual NHS ester was quenched by the addition of 15 mM ammonium bicarbonate for 30 minutes. SDA treated sample was subsequently irradiated with UV light (365 nm, UVP) for 15 minutes at room temperature.

Samples were held at a distance of 5 cm from the light source with no obstruction. Crosslinked samples were diluted with 8 M urea to a final concentration of 2 M urea. DTT and IAA were added to a final concentration of 5 mM and 10 mM, respectively, and incubated at room temperature for 20 minutes in the dark. Lys-C (FUJIFILM Wako) and Arg-C Ultra (Promega) were added to the samples at a ratio of 1:50 and incubated overnight at 37 °C with constant shaking. Peptide samples were acidified with 0.1% formic acid (FA), pooled, and concentrated to 100 µL. Enrichment of crosslinked peptides was performed by using size exclusion chromatography (SEC) with a Superdex 30 Increase column (10/300, Cytiva) on an Ӓ KTA Purifier system (Cytiva) system. Briefly, peptides were separated using isocratic flow (0.1% FA, 30% ACN, 50 mM NaCl) at a flow rate of 0.7 mL/min. Fractions were collected every 30 seconds and pooled into groups of three and desalted using Spin Tips (Pierce) for subsequent LC-MS/MS analysis.

Crosslinked peptides were then analyzed by nano-UHPLC-MS/MS (nanoElute 2, timsTOF SCP, Bruker). 1 μL of sample was directly loaded onto a PepSep column (150 μm i.d. × 250 mm, 1.5 μm C18 resin, Bruker). Samples were run at 750 nL/min over a 90 min linear gradient from 4-35% ACN with 0.1% FA. The mass spectrometer was operated in positive ion mode. For crosslink peptide identification, the MS1 scan range was 100-1700 m/z. The TIMS ramp range was 0.6 to 1.4 1/k_0_. The TIMS ramp and accumulation times locked at 75 ms. MS/MS was performed on 5 PASEF ramps per cycle. Data acquisition was performed using Bruker timsControl (version 6.1.1) and Bruker Compass HyStar (version 6.3) software.

Instrument raw files were converted into Mascot Generic Format (MGF) files using ProteoWizard ^20^. Converted raw files were loaded into Proteome Discoverer 3.1 and searched against thirteen proteins making up the pol II-Prp40 FF4-6 complex using the MS Annika 3.0 plugin ^21^. Search parameters for the search for DSSO crosslinks included carbamidomethylation-C as a fixed modification, oxidation-M, DSSO-STYK, DSSO/amidated-STYK, and DSSO/hydrolysed-STYK as variable modifications, allowing for 3 missed cleavages. Search parameters for the search for SDA crosslinks included carbamidomethylation-C as a fixed modification, oxidation-M, SDA-[Any Amino Acid], allowing for 3 missed cleavages. Precursor mass tolerance was set to 15 ppm, with MS/MS mass tolerance set to 20 ppm. Medium and high confidence FDR cutoffs (peptides and crosslinks) were set to 0.5% and 0.1%, respectively.

Results were visualized using xiVIEW ^22^.

## Acknowledgements

This work was supported by NIH R35GM145289 (R.Z.). We thank members of the Zhao lab, especially Dr. Da Cui, for their helpful feedback.

## Supplementary figure legends

**Supplementary Fig. S1.**
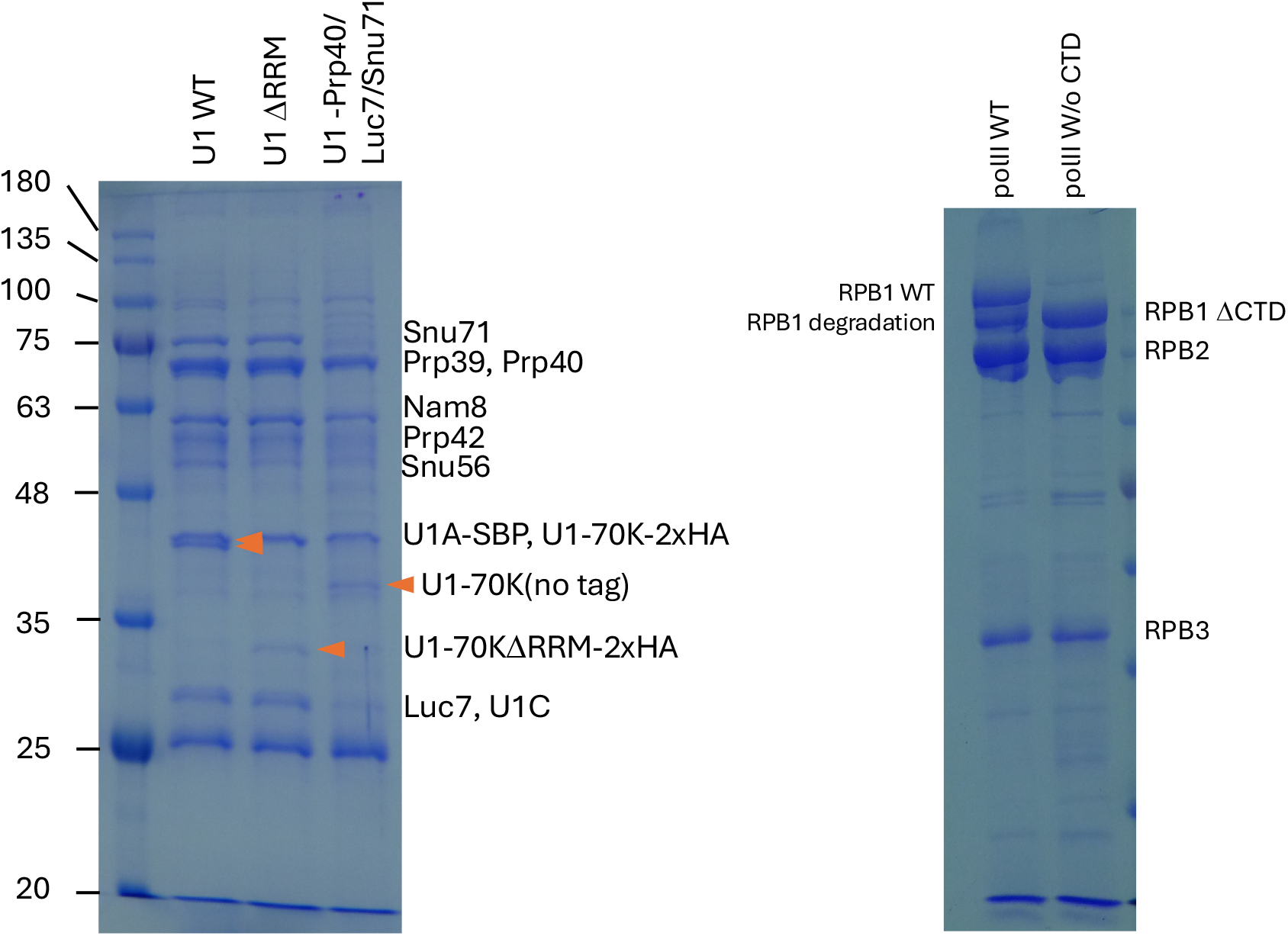
Purified U1 WT, ΔRRM, Prp40/Luc7/Snu71 depletion, pol II WT, and CTD truncation shown on SDS PAGE with Coomassie stain.

## Notes

### Competing Interest Statement

The authors have declared no competing interest.

## References

1. Herzel, L., Ottoz, D. S. M., Alpert, T. & Neugebauer, K. M. Splicing and transcription touch base: co-transcriptional spliceosome assembly and function. Nat Rev Mol Cell Biol 18, 637–650 (2017). 10.1038/nrm.2017.63

2. Phatnani, H. P. & Greenleaf, A. L. Phosphorylation and functions of the RNA polymerase II CTD. Genes Dev 20, 2922–2936 (2006). 10.1101/gad.1477006

3. Will, C. L. & Luhrmann, R. Spliceosome structure and function. Cold Spring Harbor perspectives in biology 3 (2011). 10.1101/cshperspect.a003707

4. Morris, D. P. & Greenleaf, A. L. The splicing factor, Prp40, binds the phosphorylated carboxyl-terminal domain of RNA polymerase II. J Biol Chem 275, 39935–39943 (2000). 10.1074/jbc.M004118200

5. Yu, Y. & Reed, R. FUS functions in coupling transcription to splicing by mediating an interaction between RNAP II and U1 snRNP. Proc Natl Acad Sci U S A 112, 8608–8613 (2015). 10.1073/pnas.1506282112

6. Zhang, S. et al. Structure of a transcribing RNA polymerase II-U1 snRNP complex. Science 371, 305–309 (2021). 10.1126/science.abf1870

7. Li, X. et al. CryoEM structure of Saccharomyces cerevisiae U1 snRNP offers insight into alternative splicing. Nature communications 8, 1035 (2017). 10.1038/s41467-017-01241-9

8. Jasiak, A. J. et al. Genome-associated RNA polymerase II includes the dissociable Rpb4/7 subcomplex. J Biol Chem 283, 26423–26427 (2008). 10.1074/jbc.M803237200

9. Li, Z. & Brow, D. A. A rapid assay for quantitative detection of specific RNAs. Nucleic Acids Res 21, 4645–4646 (1993).

10. Hilleren, P. J., Kao, H. Y. & Siliciano, P. G. The RRM domain is dispensable for yeast U1-70K function. Nucleic acids symposium series, 244–247 (1995).

11. Chalivendra, S. et al. Selected humanization of yeast U1 snRNP leads to global suppression of pre-mRNA splicing and mitochondrial dysfunction in the budding yeast. RNA 30, 1070–1088 (2024). 10.1261/rna.079917.123

12. Tan, C. W. et al. PRPF40A induces inclusion of exons in GC-rich regions important for human myeloid cell differentiation. Nucleic Acids Res 52, 8800–8814 (2024). 10.1093/nar/gkae557

13. Becerra, S., Montes, M., Hernandez-Munain, C. & Sune, C. Prp40 pre-mRNA processing factor 40 homolog B (PRPF40B) associates with SF1 and U2AF65 and modulates alternative pre-mRNA splicing in vivo. RNA 21, 438–457 (2015). 10.1261/rna.047258.114

14. Lorenzini, P. A. et al. Human PRPF40B regulates hundreds of alternative splicing targets and represses a hypoxia expression signature. RNA 25, 905–920 (2019). 10.1261/rna.069534.118

15. Robert, F., Blanchette, M., Maes, O., Chabot, B. & Coulombe, B. A human RNA polymerase II-containing complex associated with factors necessary for spliceosome assembly. J Biol Chem 277, 9302–9306 (2002). 10.1074/jbc.M110516200

16. Kim, J. B., Yamaguchi, Y., Wada, T., Handa, H. & Sharp, P. A. Tat-SF1 protein associates with RAP30 and human SPT5 proteins. Mol Cell Biol 19, 5960–5968 (1999). 10.1128/MCB.19.9.5960

17. Caizzi, L. et al. Efficient RNA polymerase II pause release requires U2 snRNP function. Mol Cell 81, 1920–1934 e1929 (2021). 10.1016/j.molcel.2021.02.016

18. Qiu, Z. R., Schwer, B. & Shuman, S. Two Routes to Genetic Suppression of RNA Trimethylguanosine Cap Deficiency via C-Terminal Truncation of U1 snRNP Subunit Snp1 or Overexpression of RNA Polymerase Subunit Rpo26. G3 (Bethesda) 5, 1361–1370 (2015). 10.1534/g3.115.016675

19. Agarwal, R., Schwer, B. & Shuman, S. Structure-function analysis and genetic interactions of the Luc7 subunit of the Saccharomyces cerevisiae U1 snRNP. RNA 22, 1302–1310 (2016). 10.1261/rna.056911.116

20. Chambers, M. C. et al. A cross-platform toolkit for mass spectrometry and proteomics. Nat Biotechnol 30, 918–920 (2012). 10.1038/nbt.2377

21. Birklbauer, M. J. et al. Proteome-wide non-cleavable crosslink identification with MS Annika 3.0 reveals the structure of the C. elegans Box C/D complex. Commun Chem 7, 300 (2024). 10.1038/s42004-024-01386-x

22. Combe, C. W., Graham, M., Kolbowski, L., Fischer, L. & Rappsilber, J. xiVIEW: Visualisation of Crosslinking Mass Spectrometry Data. J Mol Biol 436, 168656 (2024). 10.1016/j.jmb.2024.168656

23. UniProt, C. UniProt: the Universal Protein Knowledgebase in 2025. Nucleic Acids Res 53, D609–D617 (2025). 10.1093/nar/gkae1010

24. Madeira, F. et al. The EMBL-EBI Job Dispatcher sequence analysis tools framework in 2024. Nucleic Acids Res 52, W521–W525 (2024). 10.1093/nar/gkae241

